# A high-throughput heterologous expression platform for plant synthetic biology based on Arabidopsis suspension cells

**DOI:** 10.1101/2025.07.30.667703

**Authors:** Lukas Meile, Germán Alonso-Tolo, Zeltia Ferreiro-Eiras, Xi Jiang, Stefan Burén, Luis M. Rubio

**Author notes:** To whom correspondence should be addressed:* Luis M. Rubio: –; Lukas Meile. Other e-mail addresses: Germán Alonso-Tolo, Zeltia Ferreiro-Eiras, Xi Jiang, and Stefan Burén.

## Abstract

Efficient heterologous expression platforms are essential for plant synthetic biology, particularly for engineering complex multigene pathways. Here, we establish a high-throughput system for both transient and stable transformation of *Arabidopsis thaliana* suspension cells using plant cell pack infiltration. This method requires no specialized equipment or consumables and is compatible with several cell lines. It enables rapid generation of 100 g of transgenic cells within two weeks and allows expression of at least 6 stacked genes from a single construct. We characterized constitutive promoters for gene expression in Arabidopsis cells and validated plastid targeting peptides. A library of NifB homologs was screened for expression and solubility and several archaeal variants suitable for plant expression were identified. We further engineered stable cell lines expressing up to six genes, encoding the NifB module components NifU, NifS, FdxN, and NifB, demonstrating that the newly developed platform integrates into an established workflow for nitrogenase engineering. The platform accelerates design–build–test cycles and facilitates the production of delicate proteins that require large amounts of transgenic biomass. It thus represents a versatile and scalable tool for advancing synthetic biology and for tackling major biotechnological challenges, such as biological nitrogen fixation.

**Highlight:** We developed a fast and scalable expression platform in Arabidopsis suspension cells, enabling transient and stable multigene expression for applications in plant synthetic biology such as nitrogenase engineering.

## Introduction

Plant cell suspension cultures provide a valuable system for heterologous protein production with several advantages over whole plants (Fischer *et al*., 1999; Menges and Murray, 2004). They offer faster growth rates, simpler cultivation protocols, and reduced biological complexity due to a more uniform cell population. The homogeneous nature of suspension cultures facilitates reproducibility and scalability, which are essential for synthetic biology applications that require precise control over gene expression. Furthermore, the direct contact of all cells with the growth medium enables rapid and uniform exposure to external compounds such as expression inducers, signaling molecules, or nutrients. Together, these features make plant suspension cultures a powerful platform for accelerating design–build–test cycles in plant synthetic biology.

Several plant species have been employed for cell suspension-based expression, with tobacco (*Nicotiana tabacum*), rice (*Oryza sativa*), and carrot (*Daucus carota*) being the most widely used ones (Santos *et al*., 2016). *Arabidopsis thaliana*, although used less frequently, offers some advantages that are yet to be fully exploited. As the most extensively studied model plant, Arabidopsis provides many genetic resources, including mutant collections and a well-annotated genome (Provart *et al*., 2016). In addition, certain Arabidopsis cell lines retain the ability to form functional chloroplasts, allowing photomixotrophic growth (McCarthy *et al*., 2016; Sello *et al*., 2017) or even photoautotrophy when gradually reducing the external carbon source (Hampp *et al*., 2012). Chloroplast development in suspension cells provides opportunities to study and engineer plastid-targeted proteins and chloroplast-related processes (Li *et al*., 2021; McCarthy *et al*., 2016; Sello *et al*., 2016). Although Arabidopsis suspension cells have been used in various studies, and protocols for stable transformation and transient expression have been reported (Burén *et al*., 2011; García-León *et al*., 2018; Menges and Murray, 2004; Van Leene *et al*., 2011), throughput and fast scalability remain limited. Furthermore, as for other plant systems, integration of multi-gene modules is still not a routine application (Rizzo *et al*., 2023).

Engineering plants to fix atmospheric dinitrogen through direct transfer of prokaryotic nitrogen fixation (*nif*) genes is a key challenge of plant biotechnology (Curatti and Rubio, 2014; Oldroyd and Dixon, 2014). Nitrogenase is the prokaryotic multimeric metalloenzyme responsible for biological nitrogen fixation (BNF). Its function requires the coordinated expression of more than 10 Nif proteins, many of which are involved in the biosynthesis of nitrogenase-specific essential Fe-S cofactors (Burén *et al*., 2020). In addition to the high number of required Nif proteins, oxygen-sensitivity and low solubility of nitrogenase and other Nif proteins are major obstacles for engineering a functional plant nitrogenase (Burén and Rubio, 2018). Current strategies to engineer Nif proteins for eukaryotic expression have exploited the natural diversity of Nif proteins by systematically testing natural Nif protein variants for expression and solubility (Burén *et al*., 2019; Jiang *et al*., 2022; Jiang *et al*., 2021).

The nitrogen fixation pathway can be divided into functional modules, each mediating an independent step of nitrogenase assembly. These modules can be optimized independently, reducing complexity and providing a tractable route toward full pathway reconstruction. Beyond optimizing protein expression, successful nitrogenase engineering is also thought to require subcellular targeting, with mitochondria and plastids being the two main candidate compartments. Mitochondria offer a favorable redox environment in which oxygen is consumed by respiration, but necessarily require protein import, which might be a limiting step for some Nif proteins (Oldroyd and Dixon, 2014). Plastids, by contrast, are genetically more accessible and support high expression levels (Boehm and Bock, 2019; Scotti *et al*., 2012). Furthermore, reducing power is abundant in chloroplasts and plastid electron transport components are compatible with nitrogenase (Yang *et al*., 2017). Although oxygen production during photosynthesis poses a major obstacle (Aznar-Moreno *et al*., 2021; Eseverri *et al*., 2020b; Ivleva *et al*., 2016), strategies such as targeting non-photosynthetic plastids or temporally separating nitrogenase expression from photosynthesis have been proposed (Curatti and Rubio, 2014; Dixon *et al*., 1997). In this context, Arabidopsis suspension cultures represent a particularly versatile model. They can be cultured under heterotrophic or photomixotrophic conditions, enabling studies on how oxygen production in chloroplasts affects Nif protein stability.

Biotechnological challenges such as nitrogenase engineering highlight the need for efficient platforms that allow iterative and flexible testing of complex biosynthetic pathways in plants. Here, we describe a platform for both transient and stable transformation of Arabidopsis suspension cells and demonstrate its utility by integrating it into an established workflow for nitrogenase engineering. We characterized a suite of constitutive promoters and performed expression and solubility screenings of a NifB library, providing new promising NifB candidates suitable for plant expression. We furthermore showed that the platform supports the stacking of at least six transgenes and allows quick scalability for protein purification. By enabling rapid protein production and synthetic pathway assembly in plant cells, this simple and low-cost platform contributes to advancing experimental approaches in plant synthetic biology.

## Materials and methods

### Plant cell lines and bacterial strains

The Arabidopsis cell lines YG-1 and T87 were provided by the Riken Plant Cell Bank (Yokohama, Japan). PSB-L cells were a gift from David Alabadí (IBMCP, Valencia, Spain) and Col-0 and MM1 cells were a gift from Laszlo Bako (Umeå University, Sweden). The *Escherichia coli* strain TOP10 was used for all cloning steps. The *Agrobacterium tumefaciens* strain GV3101::pMP90 was a gift from Vicente Rubio and used for PCP infiltration if not stated otherwise.

### Arabidopsis cell suspension cultures

Col-0 cells were routinely cultured in Murashige and Skoog (MS) medium including vitamins with 3% sucrose, pH 5.7, supplemented with 2,4-dichlorophenoxyacetic acid (2,4-D, 0.24 µg/ml) and kinetin (0.014 µg/ml) with weekly passages of 7 ml culture into 40 ml of fresh medium in 250-ml flasks. MM1 and PSB-L cells were routinely cultured in 1x MS Basal Salts medium with 3% sucrose, pH 5.7, supplemented with thiamine (0.4 µg/ml), myo-inositol (0.1 mg/ml), 1-naphthaleneacetic acid (NAA, 0.5 µg/ml) and kinetin (0.05 µg/ml) with weekly passages of 2 ml culture into 48 ml of fresh medium and 10 ml culture into 40 ml of fresh medium, respectively. YG-1 cells were cultured in 1x MS Basal Salts medium with 200 mg/L KH_2_PO_4_, 3% sucrose, pH 5.8, supplemented with thiamine (0.4 µg/ml), myo-inositol (0.1 mg/ml), and 2,4-D (0.2 µg/ml) with weekly passages of 5 ml sedimented cells into 40 ml of fresh medium. T87 cells were cultured in MS medium including vitamins with 1.5% sucrose, pH 5.7, supplemented with NAA (0.224 µg/ml) with biweekly passages of 1.2 ml of culture into 40 ml fresh medium. Col-0 and YG-1 cells were routinely cultured at 25°C without light and MM1, PSB-L, and T87 cells were cultured at 22°C under a 16-h photoperiod.

### Transient and stable plant cell transformation

The methods for transient and stable transformation of Arabidopsis suspension cells are summarized in Fig. 1. Precultures for transient expression assays were prepared as described above and grown for 3-7 days before transformation. Transient expression assays were based on the plant cell pack (PCP) infiltration method (Rademacher *et al*., 2019). For PCP infiltration of Col-0, the medium was first removed by pressing the tip of a serological pipette against the flask or tube bottom followed by slow aspiration, similarly as described (Cortese *et al*., 2021). The resulting compacted cells were then resuspended in Paul’s Medium (Buschmann, 2016) to a density that barely allowed pipetting of the suspension with wide-bore tips. For small PCPs of approximately 200 mg (fresh weight), 500 µl of cell suspension was loaded onto the top part of 1000-µl filter pipette tips. The excess liquid was removed by vacuum for approximately 10 s, resulting in PCP formation. *A. tumefaciens* suspensions were prepared from 1-day-old precultures as described before (Gengenbach *et al*., 2020), but without acetosyringone, at OD_600nm_ 0.4 if not stated otherwise. PCPs were infiltrated with 300 µl of *A. tumefaciens* suspension and incubated at 25°C for 1 h before removing the excess liquid by vacuum for approximately 30 s. The infiltrated PCPs were incubated inside a tip box equipped with sterile wet paper to maintain high humidity for 2-4 days at 25°C in the dark, unless indicated otherwise.

**Fig 1:**
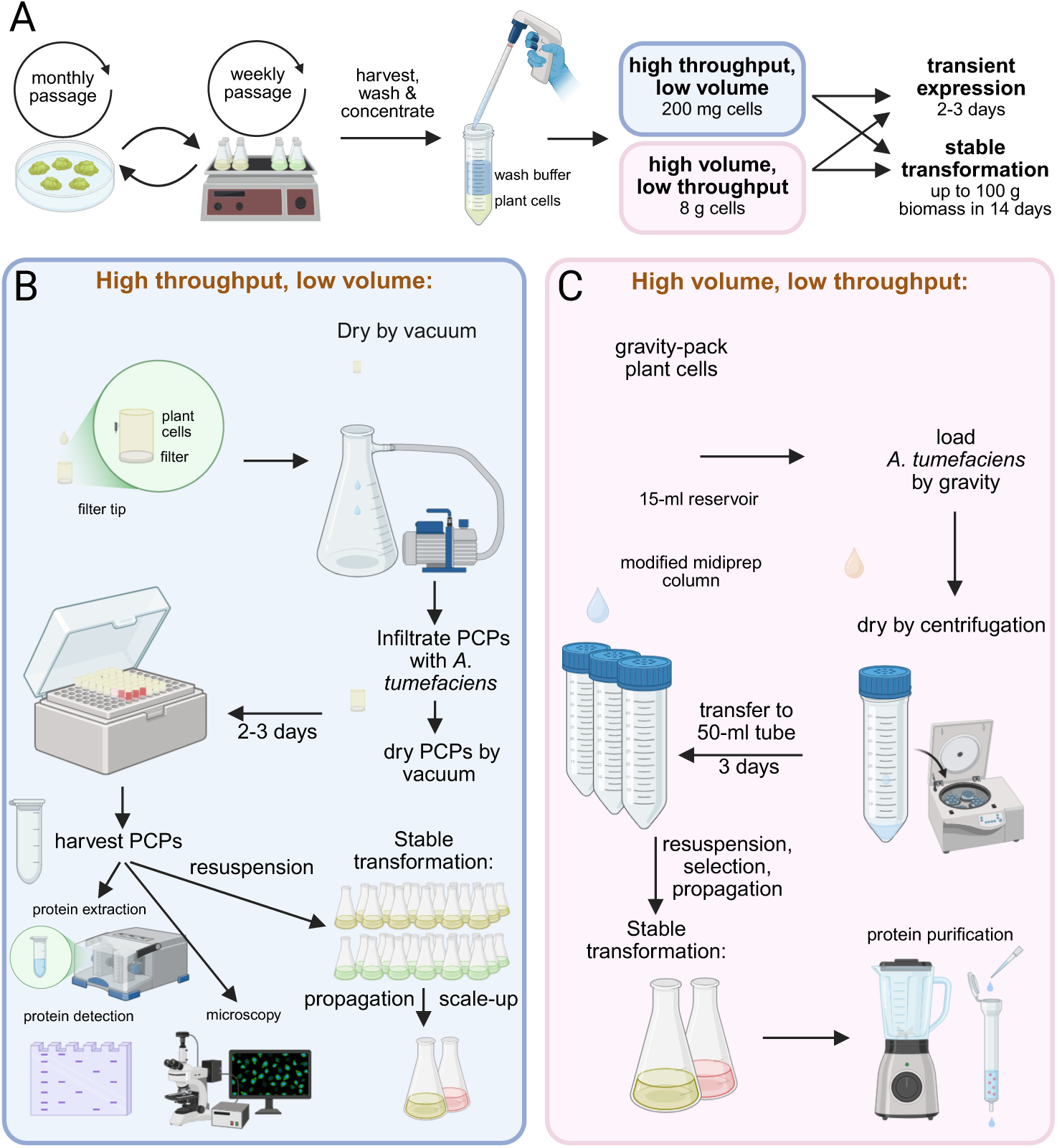
Workflow and applications of plant cell pack (PCP) infiltration in Arabidopsis suspension cells for stable and transient transformation at different scales. (A) Plant cells are maintained by weekly or monthly passaging in liquid or solid MS medium, respectively. Washed plant cells grown in liquid medium under darkness serve as starting material for both transient and stable transformations. 200-mg PCPs allow high throughput, whereas 8-g PCPs allow the production of up to 100 g (fresh weight) of transgenic cells in 2 weeks. (B) For high-throughput applications, a dense suspension of washed cells is loaded on top of a pipette filter tip. The excess liquid is then removed by vacuum and the semi-dry PCP is infiltrated with an *A. tumefaciens* suspension. After 60 min, the excess liquid is removed and the PCPs show transgene expression after 2-3 days. Individual PCPs can be conveniently transferred to microcentrifuge tubes downstream applications such as protein extraction, microscopy, or the generation of stably transformed cells. (C) For high-volume applications such as the purification of heterologously produced proteins, washed suspension cells are loaded onto midi-prep columns and excess liquid is removed by gravity or gentle vacuum. *A. tumefaciens* is loaded by gravity and, after 60 min, excess liquid and bacteria are removed by gentle centrifugation and PCPs are transferred to 50-ml centrifugation tubes. After 2-3 days, transformed PCPs are resuspended in fresh medium to start a liquid culture. Created in https://BioRender.com.

For small-scale stable transformations of around 200 mg of starting biomass, PCPs were prepared the same way as for transient expression assays. For larger PCPs of approximately 8 g, cell suspensions were prepared similarly but 20 ml was loaded onto a Zymo-Spin V-PS midiprep column assembly (with the DNA-binding matrix removed) with a 15-ml Reservoir-X (Zymo Research, CA, USA). The excess medium was removed by gravity and 20 ml of *A. tumefaciens* suspension was passed through the packed cells by gravity. The excess liquid was removed by centrifugation in a swing-bucket rotor at 200 g for 2 min. The compacted, semi-dry PCPs were then transferred to a 50-ml centrifugation tube with a partially closed lid, sealed with Micropore tape (3M Health Care, USA) and incubated in a moist box. After 2-4 days, small PCPs were gently resuspended in 4 ml of cultivation medium described above containing carbenicillin (250 µg/ml) and vancomycin (165 µg/ml) to select against *A. tumefaciens*. For cell lines Col-0 and YG1, selection with hygromycin was applied at this point (20 µg/ml), whereas for MM1 hygromycin was only applied 2-3 days after resuspension. For large PCPs, cells were resuspended in 40 ml of growth medium, agitated on an orbital shaker at 100 rpm for 30 min before replacing the medium and supplementing with antibiotics as for small PCPs.

### Plasmids, molecular cloning, gene library design and codon optimization

Molecular cloning was performed using the Modular Cloning (MoClo) standard (Weber *et al*., 2011; Werner *et al*., 2012) as described before (Eseverri *et al*., 2020b). For domestication of Level 0 parts, the Phusion Hot Start II DNA Polymerase (Thermo Fisher) was used with the primers and templates listed in Table S1. The sequence of all generated constructs used in this study was verified using a Sanger sequencing service (Macrogen, Madrid, Spain) or an Oxford Nanopore whole-plasmid sequencing service (Plasmidsaurus, Inc., Eugene, USA). All constructs and the individual parts used for their assembly are listed in Table S2. The MoClo Plant Parts Kit and pUAP1 were a gift from Nicola Patron (Addgene kit #1000000047, and plasmid #63674, respectively) and the MoClo Toolkit was a gift from Sylvestre Marillonnet (Addgene kit #1000000044). 35S:RUBY was a gift from Yunde Zhao (Addgene plasmid #160908) and pTHsp18.2 (GB0035) was a gift from Diego Orzaez (Addgene plasmid #68186). The *nifB* library was designed by keyword search in the Uniprot database (Bateman *et al*., 2021). Archaeal NifB homologs were identified by discarding protein sequences lacking the six described NifB characteristic motifs (Arragain *et al*., 2017), resulting in 321 candidate archaeal NifB homologs. Because Nif proteins from thermophilic organisms have been successfully expressed in eukaryotes before (Burén *et al*., 2017; Jiang *et al*., 2022), NifB homologs from thermophilic sources were considered as high-priority candidates. To identify them, NCBI taxon IDs associated with the Uniprot search results were used to obtain information on growth temperature and habitats in the Bac*Dive*, JGI IMG/D and GOLD databases (Chen *et al*., 2021; Mukherjee *et al*., 2025; Schober *et al*., 2025). Furthermore, homologs from known diazotrophs were considered good candidates. Diverse source organisms were also considered by not including more than two homologs from the same taxonomic order. The workflow for the library design is summarized in Fig. S1. To improve expression in Arabidopsis, all *nifB* sequences as well as the sequences of *nifU*, *nifS*, *fdxN*, and the TS-tag were codon optimized using the DNA Chisel tool (Zulkower and Rosser, 2020). The optimization constraints were high GC and GC3 contents of at least 50 and 69%, respectively and avoiding BpiI, BsaI, and BsmBI restriction sites. The optimization objectives included codon optimization for Arabidopsis, avoiding rare codons (minimum frequency = 0.25), avoiding hairpins (stem size = 15, window = 200) as well as avoiding the instability motifs considered by previous work (Eseverri *et al*., 2020b). The optimized sequences flanked by BsaI sites for cloning (Table S3) were ordered from GenScript (USA).

### Luciferase reporter assay

Promoter activities were measured as previously reported (Li *et al*., 2017) with modifications. PCPs were lysed in a buffer containing 100 mM Tris-HCl (pH 8.6), 200 mM NaCl, 10% glycerol, 0.1% protease inhibitor cocktail (P9599, Sigma-Aldrich) with 4:1 (v/w) buffer/biomass ratio by metal bead (7 mm) beating in a Qiagen Retsch MM300 TissueLyser for 60 s. Ten µl cell lysate was mixed with 100 µl luciferase buffer containing 25 mM glycyl-glycine (pH 8), 14 mM K_2_HPO_4_-KH_2_PO_4_ (pH 8), 4 mM EGTA, 2 mM ATP, 1 mM DTT, 15 mM MgSO_4_, 150 µg/ml D-luciferin in a white 96-well plate. Luminescence was measured with an integration time of 2 s in a Varioskan LUX plate reader (Thermo Scientific). For normalization of luciferase levels, every construct contained the same GUS reporter cassette. GUS activity was measured in the same device by mixing 10 µl cell lysate with 50 µl buffer containing 10 mM Tris-HCl (pH 8.0), 1 mM 4-methylumbelliferyl-β-D-glucuronide, and 2 mM MgCl_2_ in a black 96-well plate. The reaction was incubated at 37°C for 30 min with 100-ms measurements in 15-s intervals with an excitation wavelength of 365 nm with a bandwidth of 12 nm and an emission wavelength of 455 nm. The GUS activity was determined by calculating the change in fluorescence over time in the linear phase. The promoter activity was quantified by dividing the luminescence intensity by the GUS activity and normalized to the strongest promoter of each run.

### Confocal microscopy

PCPs of the cell line Col-0 were infiltrated with *A. tumefaciens* harbouring the plasmids 1608-1614 (Table S2) as described above and harvested after 2 days by resuspension in growth medium. Confocal laser scanning microscopy was performed with a Zeiss LSM 880 microscope equipped with a Plan-Apochromat 40X/1.2 water-immersion objective and 488-nm and 561-nm lasers. The detection ranges were 493-574 nm for GFP and 578-696 nm for mCherry. Images were acquired using the ZEN 2.6 Black software (Zeiss) and processed with the Fiji software (Schindelin *et al*., 2012); modifications included cropping, maximum-intensity projections, and adjustments of brightness and contrast.

### Protein extraction, immunoblotting and purification

Protein extraction for immunoblotting was performed similarly as described (Burén *et al*., 2017) with modifications, if not stated otherwise. PCPs were added to a 2-ml Eppendorf tube and subjected to 4 freeze-thaw cycles with liquid N_2_. Lysis buffer, containing 100 mM Tris-HCl (pH 8), 150 mM NaCl, 10 mM MgCl_2_, 0.2% NP-40, 5% glycerol, 5 mM β-mercaptoethanol, 5 mM EDTA, and 0.5% protease inhibitor cocktail (Sigma P9599; added immediately before use), was added at a ratio of 2:1 (v/w) with a 7-mm diameter steel ball and the cells were homogenized in a Qiagen Retsch MM300 TissueLyser for 60 s, followed by a 15-min incubation at 4°C in a tube rotator. The resulting homogenate was mixed at a ratio of 1:1 with 2x Laemmli buffer (125 mM Tris-HCl pH 6.8, 4% SDS, 20% (v/v) glycerol, 10% β-mercaptoethanol, 0.005% bromophenol blue) and heated for 10 min at 98°C, yielding the total cell extract. For the soluble extract, the cell homogenate was centrifuged at 20,000 g for 15 min and the supernatant containing the soluble proteins was treated with 2x Laemmli buffer as described for the total extract. The heat-denatured protein samples were centrifuged at 20,000 for 10 min before SDS-PAGE electrophoresis and immunoblotting. After blotting, membranes were stained with a Ponceau S solution (0.2% in 5% acetic acid) as transfer control. The Twin-Strep (TS) tag was detected with a monoclonal mouse StrepII antibody (GenScript A01732, 1:5,000), NifU and NifS with rabbit polyclonal anti-NifU and anti-NifS antibodies (Rubio lab, 1:20,000), the HA-tag with a rat monoclonal antibody (Roche 12013819001, 1:5,000), GFP with a mouse monoclonal antibody (Santa Cruz sc-9996, 1:2,000), and RbcL with a rabbit polyclonal antibody (Agrisera AS03 037, 1:5,000). Secondary HRP-conjugated antibodies were used in 2% milk powder in TBS-T diluted 1:20,000 (anti-mouse and anti-rabbit) or 1:5,000 (anti-rat). NifB purifications were performed from Col-0 cells grown in the dark. Cells were transformed with plasmid pN2LM282 as described above and scaled up to four 250-ml shake flasks with 60 ml culture each. To collect cells, the grown transformant cultures were bubbled with pure N_2_ and then moved to an anaerobic chamber, where the following steps were performed. Cells were harvested by filtration through a ZymoPURE Syringe Filter-X (Zymo Research, CA, USA) followed by drying with tissue paper and snap-freezing and storage in N_2_. Cells were homogenized in a blender (Oster Classic 4655) for 5 min at maximum speed with lysis buffer (100 mM Tris-HCl [pH 8.6], 200 mM NaCl, 10% glycerol, 5 mM β-mercaptoethanol, 2 mM sodium dithionite, 1 mM phenylmethylsulfonyl fluoride, 1 mg/ml leupeptin, 5 mg/ml DNase I) with a buffer/biomass ratio of 2:1. The homogenate was then filtered through 2 layers of miracloth. Lysate clearing, NifB purification and concentration was performed as described (Jiang *et al*., 2022), but with a 5-ml Strep-Tactin®XT 4Flow® column and a peristaltic pump setup. Protein purity was assessed by SDS-PAGE analysis followed by peptide mass fingerprinting of individual bands excised from the SDS-PAGE gel. Mass spectrometry analysis was performed by a service of the Proteomic Unit at Universidad Complutense de Madrid.

## Results

### A transient expression system for Arabidopsis suspension cells

We first sought to improve the throughput of the *A. tumefaciens*-mediated transient expression workflow for Arabidopsis suspension cells. The method was based on a recently developed approach of plant cell pack (PCP) infiltration, which was originally established for tobacco BY-2 cells (Rademacher *et al*., 2019). For easy and individual handling of PCPs, they were cast into the top part of filter pipette tips and the RUBY (He *et al*., 2020) and GFP reporters were used to assess successful transformation. We first assessed four different *A. tumefaciens* strains for their suitability for PCP infiltration with different Arabidopsis cell lines and found that the strain GV3101::pMP90 led to RUBY expression in the cell line Col-0 (Fig. 2A) and YG-1 (Fig. S2A), but not in the cell lines MM1, PSB-L, and T87 (Fig. 2A). Strain AGL-1 led to RUBY expression in Col-0 but not in the other tested cell lines MM1, PSB-L, and T87 (Fig. 2A). Similarly, strain EHA105 led to RUBY expression only in Col-0 but not in the other tested cells MM1 and PSB-L (Fig. S2A). Finally, PCPs infiltrated with the strain LBA4404 never produced visible RUBY signal (Fig. 2A). The culture method of *A. tumefaciens* did not affect infiltration success, as cultures grown in liquid or solid media both led to RUBY expression (Fig. S2B). Since both Col-0 and YG-1 were routinely cultured in darkness and the other cell lines under a 16-h photoperiod, we hypothesized that cultivation under light might hamper transformation. Col-0 and MM1 cells grown either in darkness or under a 16-h photoperiod were then compared for their receptiveness to PCP infiltration. MM1 cells grown under darkness were more amenable to PCP infiltration; however, only small PCPs cast from 150 µl cell suspension but not from 500 µl consistently yielded high transformation efficiency (Fig. 2A & D). A similar effect of light conditions on transformation efficiency was observed for Col-0 cells, for which transformation was abolished when cultured under a 16-h photoperiod (Fig. 2C).

**Fig 2:**
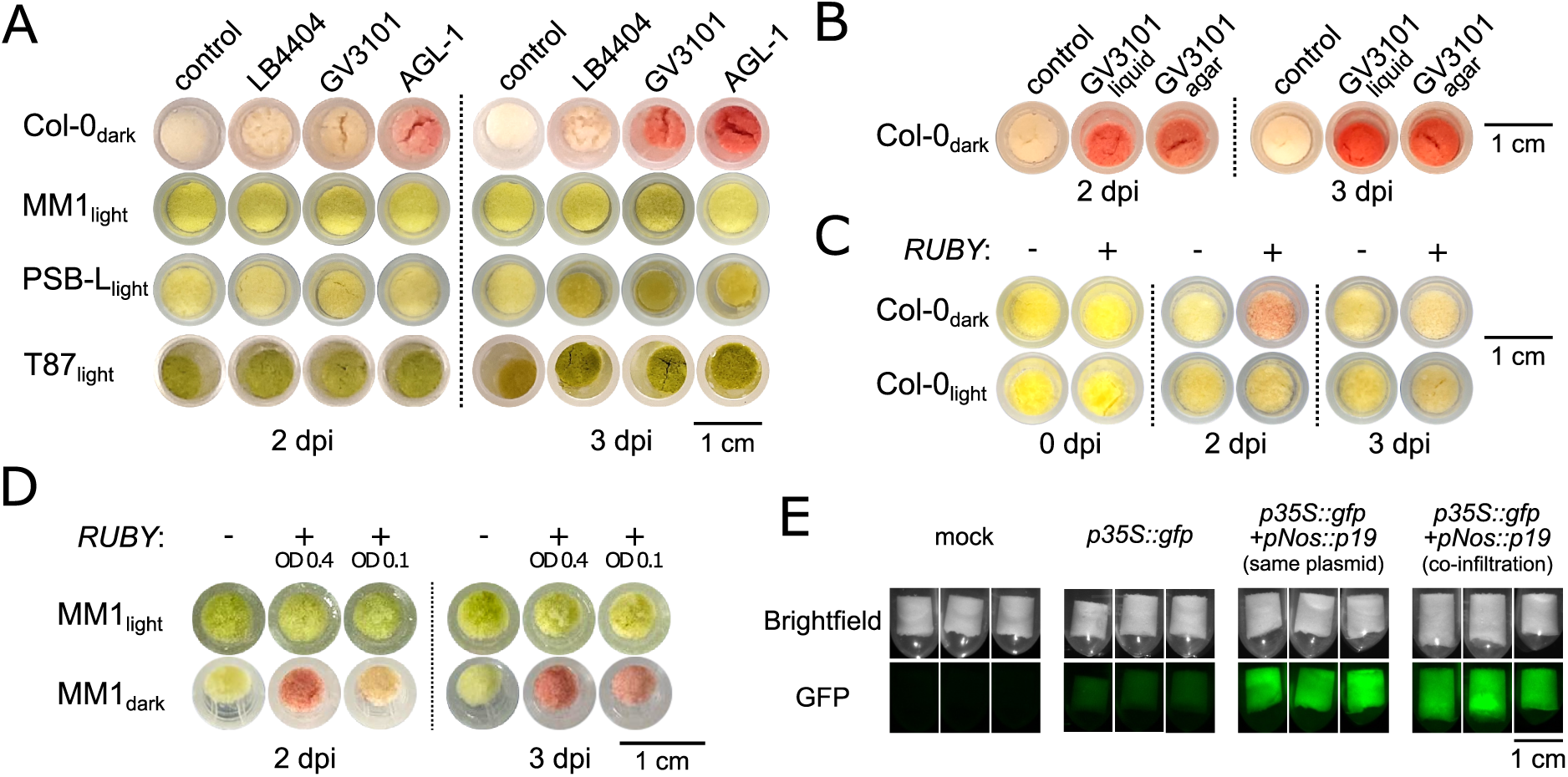
Optimization of transient expression in Arabidopsis suspension cells by plant cell pack (PCP) infiltration. (A) PCPs of four different Arabidopsis cell lines infiltrated with three different *A. tumefaciens* strains harboring a plasmid with the RUBY reporter. Pictures were taken 2 and 3 days post infiltration (dpi). PCPs treated with the infiltration buffer only were included as negative control. (B) PCPs of the Arabidopsis cell line Col-0 infiltrated with the *A. tumefaciens* strain GV3101::pMP90 carrying the RUBY cassette prepared from liquid or solid culture medium. (C) PCPs from the cell line Col-0 grown either under constant darkness or a 16-h photoperiod and infiltrated with *RUBY*-carrying *A. tumefaciens* strain GV3101::pMP90 (+) or with infiltration buffer only (-). (D) PCP infiltration of cell line MM1 grown either under darkness or a 16-h photoperiod with RUBY-carrying *A. tumefaciens* strain GV3101::pMP90 prepared at different optical densities (+) or with infiltration buffer only (-). (E) Co-expression of the suppressor of silencing P19 driven by the *Nos* promoter and GFP driven by the *35S* promoter in PCPs of cell line Col-0. The *p19* and *gfp* genes were either present on the same or on different plasmids. Three technical replicates are shown.

Next, we tested whether Arabidopsis PCP infiltration can be improved by the co-expression of the inhibitor of silencing p19. Co-expression of GFP and p19 in Col-0 PCPs was performed, either using a single *A. tumefaciens* strain harbouring both genes on the same plasmid or using two co-infiltrated strains, each harbouring one of the genes. To avoid the detection of GFP expressed in *A. tumefaciens* cells, which has been described before (Buschmann, 2016), a *gfp* version containing the *DEM1* intron (Christie *et al*., 2011) was deployed. The p19 improved GFP expression irrespectively of whether *gfp* and *p19* were present on the same plasmid or co-infiltrated (Fig. 2E).

### PCP infiltration is a fast method for generating stably transformed plant cells

We hypothesized that infiltrated PCPs could serve as an efficient source for stably transformed cell cultures. This hypothesis was tested with the *A. tumefaciens* strain GV3101::pMP90 because this strain has been successfully used for stable transformation of Arabidopsis suspension cells (Burén *et al*., 2011; García-León *et al*., 2018). When resuspended in fresh culture medium amended with carbenicillin and vancomycin to eliminate *A. tumefaciens*, PCPs carrying a hygromycin selection cassette quickly formed a transgenic suspension culture under hygromycin selection, as demonstrated by expression of the RUBY reporter (Fig. S2C-E). For the cell lines Col-0 and MM1, transformation was possible with small PCPs of approximately 200 mg and 60 mg, respectively, as well as with large PCPs of approximately 8 g. The transformation of large PCPs followed by hygromycin selection and scaling up the culture allowed the production of nearly 100 g (fresh weight) of transformed Col-0 cells within 13 days after PCP infiltration (Fig. S2D). Similarly to what was observed for transient expression, the formation of transgenic lines from PCPs prepared from the cell line MM1 was only consistently possible if the cells were grown under darkness before infiltration (Fig. S2E). Furthermore, 2 days of co-culture with *A. tumefaciens* yielded more consistent transformations than 3 days and an optical density of the applied *A. tumefaciens* of 0.1 favoured transformation compared to a higher density of 0.4 (Fig. S2E).

### Characterization of a promoter set for heterologous expression in plant cells

To expand the molecular toolkit for synthetic biology applications including heterologous expression from multigene constructs, we characterized 12 promoters for constitutive gene expression in Arabidopsis suspension cells. A luciferase-based reporter assay was used to measure promoter activity in infiltrated PCPs. The cassava vein mosaic virus (CsVMV) promoter led to the highest luciferase expression and the activity of the other promoters is therefore shown as percentage of the CsVMV activity (Fig. 3). The widely used cauliflower mosaic virus *35S* (CMV35S) promoter exhibited similar activity (96±16%), followed by the Arabidopsis ubitquitin-10 (*AtUbq10)* promoter (73±3%) and the *A. tumefaciens* mannopine synthase (*AtuMas*) promoter (30±12%). The tomato histone H4 (*SlH4*) promoter and the *A. tumefaciens* nopaline synthase (*AtuNos*) promoters were considerably weaker (3±1% and 3±3%, respectively; Fig. 3). The activities of the 6 weakest promoters were between 0.6±0.2% and 0.3±0.2%. These findings suggest that the CsVMV, the CVM35S, the *AtUbq10*, and the *AtuMas* promoters are most suitable for heterologous expression of proteins to be purified and biochemically characterized in Arabidopsis suspension cells. The weaker promoters can be used to drive the expression of accessory proteins that are required at lower quantities.

**Fig 3:**
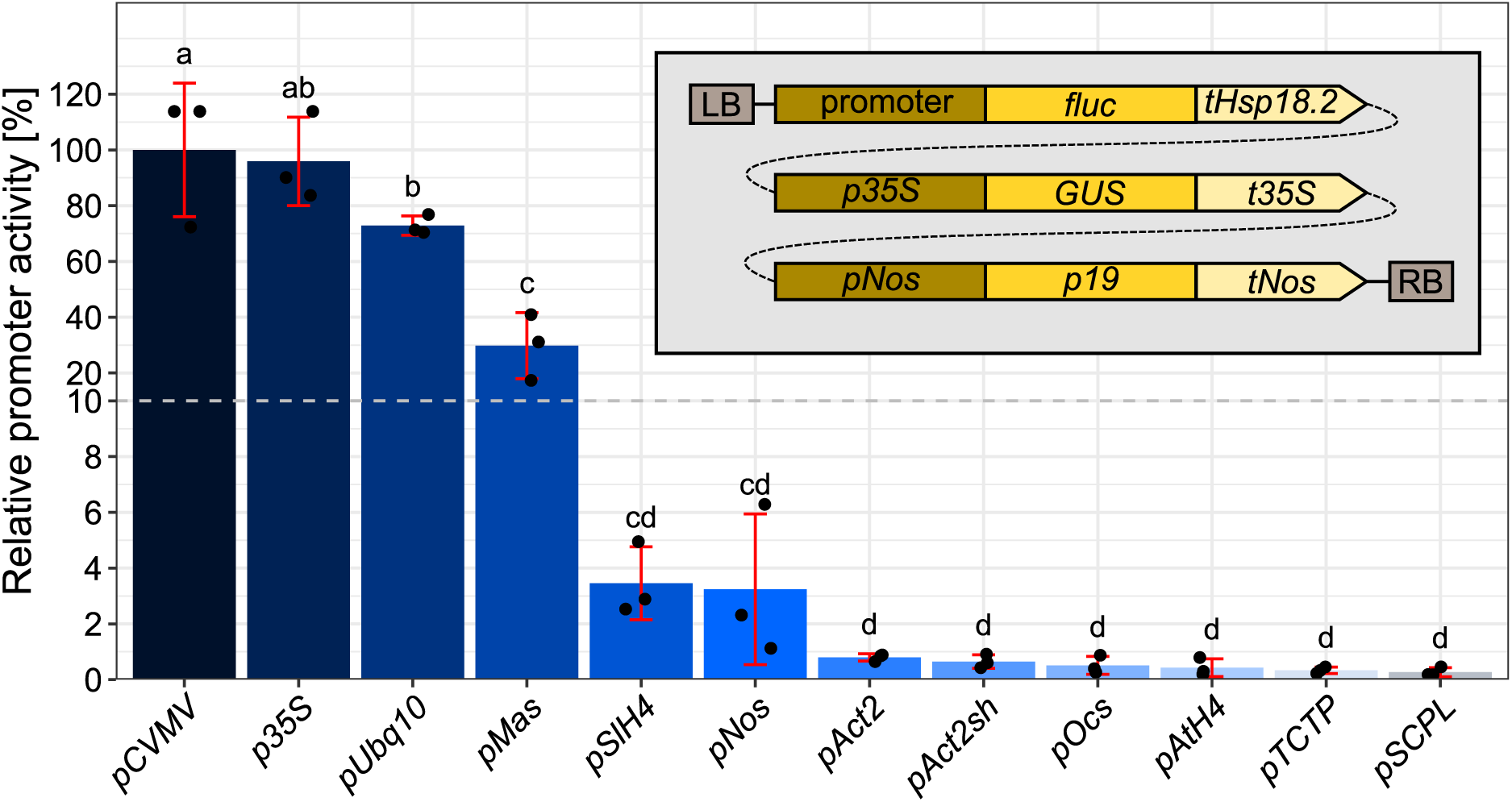
Various constitutive promoters of different strength allow transgene expression in Arabidopsis suspension cells. Relative promoter activity as determined by LUC reporter assays. Different constitutive promoters driving expression the firefly luciferase (*fluc*) gene were co-transformed with a GUS reporter cassette (*p35S::GUS::t35S*) for normalization and the suppressor of silencing *p19* from a single T-DNA (grey box; not to scale). Promoter activity was defined as luminescence divided by GUS activity. Bars show the average of 3 biological replicates. The y-axis is segmented with two different scales. Individual replicates are shown as black dots and error bars represent the standard deviation. Letters represent statistical groups according to one-way ANOVA and Tukey’s test. LB: left T-DNA border; RB: right T-DNA border.

### Solubility screening of plastid targeted proteins

We next sought to apply the newly established method for transient and stable transformation of Arabidopsis suspension cells and the promoters validated for this system to the engineering of BNF through heterologous expression of prokaryotic nitrogen fixation (Nif) proteins. We aimed to identify new NifB protein variants that can be used to engineer BNF in plants by performing a solubility screen of naturally occurring NifB variants. Since NifB function and potentially also solubility depend on the accessory proteins NifU, NifS and FdxN, we first generated a stable cell line expressing those proteins to be used as a background line for the NifB solubility screening. We therefore first tested a subset of the previously characterized promoters and previously described chloroplast targeting peptides (CTPs) for expression and plastid localization of NifU, NifS, and FdxN. NifX was co-expressed from the same construct to test its solubility due to its function in nitrogenase co-factor biosynthesis downstream of NifB. We found that the Arabidopsis *ubiquitin-10* promoter, the tomato histone *H4* promoter and the Cassava Vein Mosaic Virus (CsVMV) promoter led to the accumulation of NifU, NifS, and FdxN, respectively, in the soluble fraction of protein extracts (Fig. 4A), whereas NifX was not soluble (Fig. S3). Since the latter protein is not needed for NifB activity, *nifX* was excluded from later constructs. The Arabidopsis CTPs from the TOCC, CAB6 and GLTB2 proteins allowed plastid localization, as assessed by the presence of a single band in immunoblots, suggesting the absence of full-length proteins with uncleaved CTPs (Fig. 4A), and by the co-localization of GFP-fusion proteins with an established plastid marker protein, as determined by confocal microscopy (Fig. 4B and Fig. S4). In a cell line that stably expressed NifU, NifS, and FdxN, a library of 15 different *BCCP1::TS::nifB* homologs was tested in transient expression assays (Fig. 5A), employing a TS-tag sequence that contains an intron to avoid false positive detection resulting from misexpression in *A. tumefaciens*. Since archaeal Nif variants have been successfully expressed in eukaryotes before, we tested 13 *nifB* homologs from archaea and only two from bacteria. Two archaeal homologs originated from uncultivated samples of extreme environments, the others are from thermophilic and/or known diazotrophic archaea (Fig. 5B, Fig. S1, Table S4). Comparing total and soluble extracts of *nifB*-infiltrated PCPs led to the accumulation of soluble NifB protein for 11 out of 15 tested NifB homologs (Fig. 5C, Fig. S5). None of the two tested bacterial NifB homologs were soluble in plant cells and only two of the tested archaeal homologs were insoluble.

**Fig 4:**
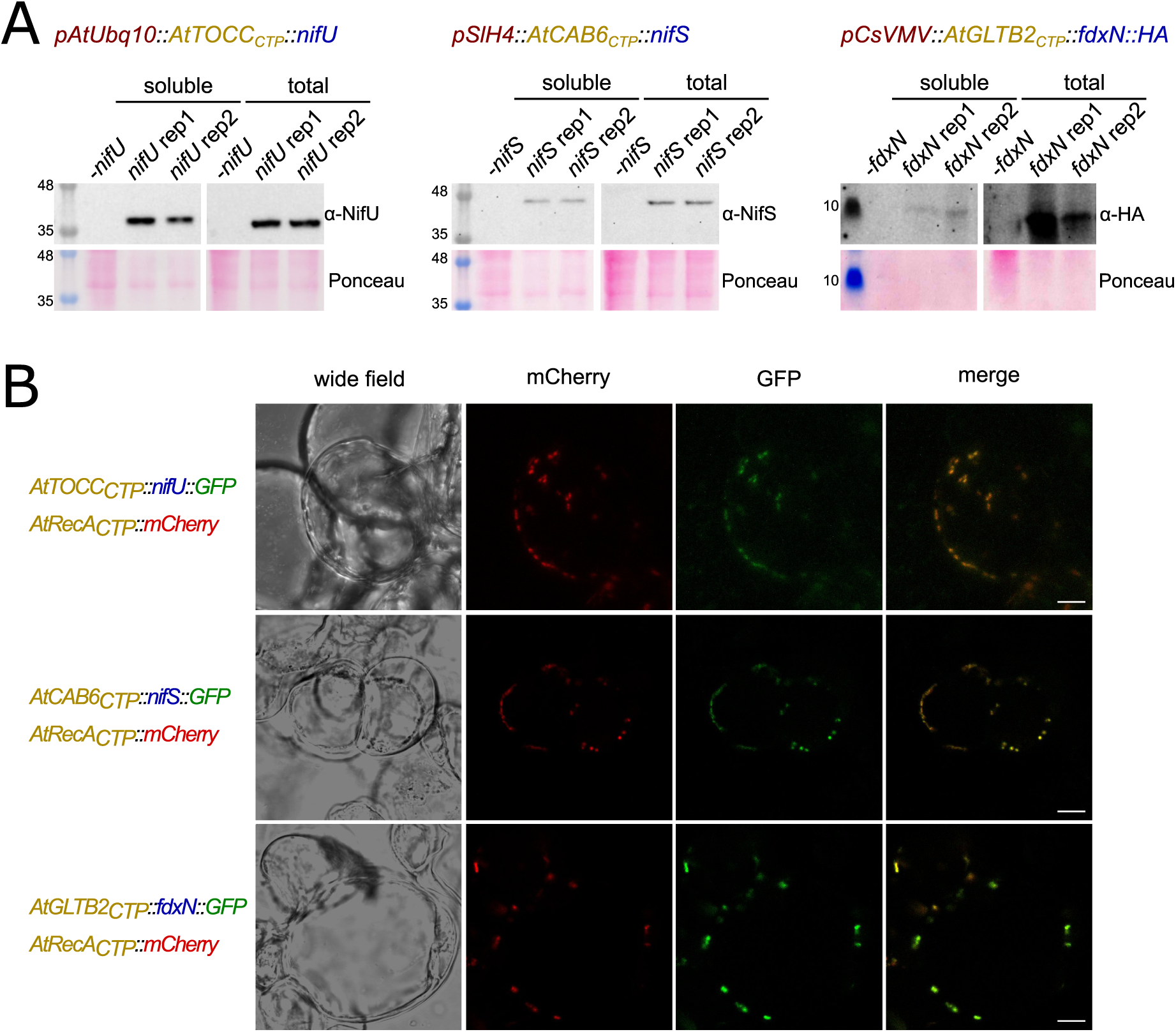
Soluble proteins of the NifB functional module can be expressed and targeted to plastids in Arabidopsis suspension cells. (A) Western blots of total and soluble plant cell extracts transformed with *nifU*, *nifS*, and *fdxN* driven by the Arabidopsis *ubiquitin-10* promoter, the tomato histone H4 promoter, and the Cassava Vein Mosaic Virus (CsVMV) promoter, respectively. The proteins were fused to minimal versions of the chloroplast-targeting peptides (CTPs) of the Arabidopsis proteins TOCC, CAB6, and GLTB2, respectively (Eseverri *et al*., 2020a). FdxN was fused to an HA-tag for detection, whereas NifU and NifS were untagged. Col-0 cells were transformed with plasmid pN2LM165 (rep1) and plasmid pN2LM166 (rep2) and proteins extracted with a buffer containing 100 mM Tris-HCl (pH8), 150mM NaCl, 10% glycerol and 0.5% protease inhibitor cocktail. (B) Detection of GFP signal in Arabidopsis cells transformed with *nifU*, *nifS* and *fdxN* fused to *gfp* and the TOCC, CAB6 and GLTB2 CTPs, respectively, by confocal laser scanning microscopy. The mCherry reporter fused to the RecA CTP was used as a control for plastid localization. Merged pictures of the mCherry and GFP channels are also shown. Scale bars: 10 µm.

**Fig 5:**
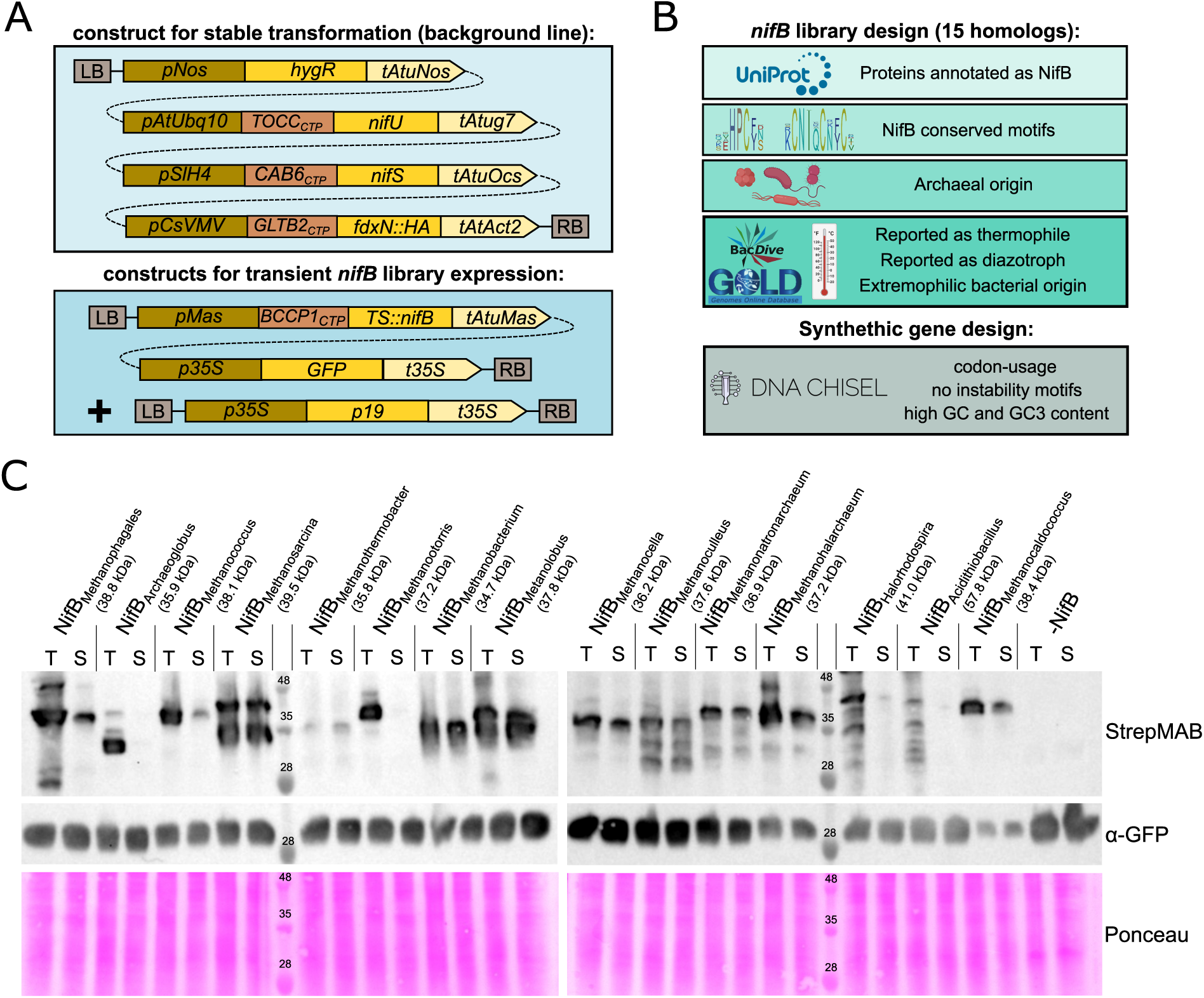
Solubility screening identified new archaeal NifB protein variants suitable for plant expression. (A) Construct and expression design for the NifB solubility screening. A stably transformed background line expressing *nifU*, *nifS*, and *fdxN::HA* was generated first. In this background line, the different *nifB* homologs fused to the BCCP1 chloroplast targeting peptide (Eseverri *et al*., 2020b) were transiently co-expressed with *gfp* as a transformation control and the suppressor of silencing p19. (B) Workflow overview for the *nifB* library and synthetic gene design. NifB variants annotated by Uniprot were filtered for known NifB protein motifs (Arragain *et al*., 2017) and for archaeal origin. Public databases such as Bac*Dive* (Schober *et al*., 2025) and information on isolation sources were used to select 8 NifB homologs from thermophilic organisms. Additionally, four homologs from known diazotrophic archaea, one from an uncultured archaeon, and two from extremophilic bacteria were included. The synthetic gene design was performed using the DNA Chisel program (Zulkower and Rosser, 2020), focusing on matching the codon usage to Arabidopsis and obtaining high GC and GC3 content as well as avoiding known mRNA instability motifs. (C) Western blots of total (T) and soluble (S) plant cell extracts. NifB variants were fused to the Twin-Strep (TS) tag for detection with a Strep monoclonal antibody (StrepMAB). A control with the same expression vector but without *nifB* was included (-NifB). LB: left T-DNA border; RB: right T-DNA border; CTP: chloroplast targeting peptide; *hygR*: hygromycin selection marker.

### Gene stacking and protein purification

To test the capacity of this method for stable expression of complex multi-gene constructs, we generated Arabidopsis suspension cell lines carrying a single T-DNA with six genes: a hygromycin resistance marker (*hygR*) for selection, *gfp* as a transformation control, and the four gene components of the NifB module (*nifU*, *nifS*, *fdxN*, and *nifB*). The construct design is shown in Fig. 6A. All four proteins were detected in the soluble fraction of protein extracts from stably transformed cells by immunoblotting (Fig. 6B), confirming successful expression. To further validate expression and protein integrity, we performed affinity purification using a TS-tag fused to NifB. Coomassie-stained gels of purification fractions revealed bands at the expected size alongside some nonspecific background (Fig. 6C), and a Western blot confirmed that NifB was specifically enriched during the purification (Fig. 6D). Additional immunoblots for GFP and the plastid-localized RbcL protein confirmed the release of both cytosolic and plastid contents during tissue homogenization. Mass spectrometry confirmed the identity of the purified protein of the expected mass (39.5 kDa) as TS::NifB (Fig. S6). Together, these results demonstrate that our platform supports large multi-gene constructs and simultaneous expression of several plastid-targeted proteins in plant cells.

**Fig 6:**
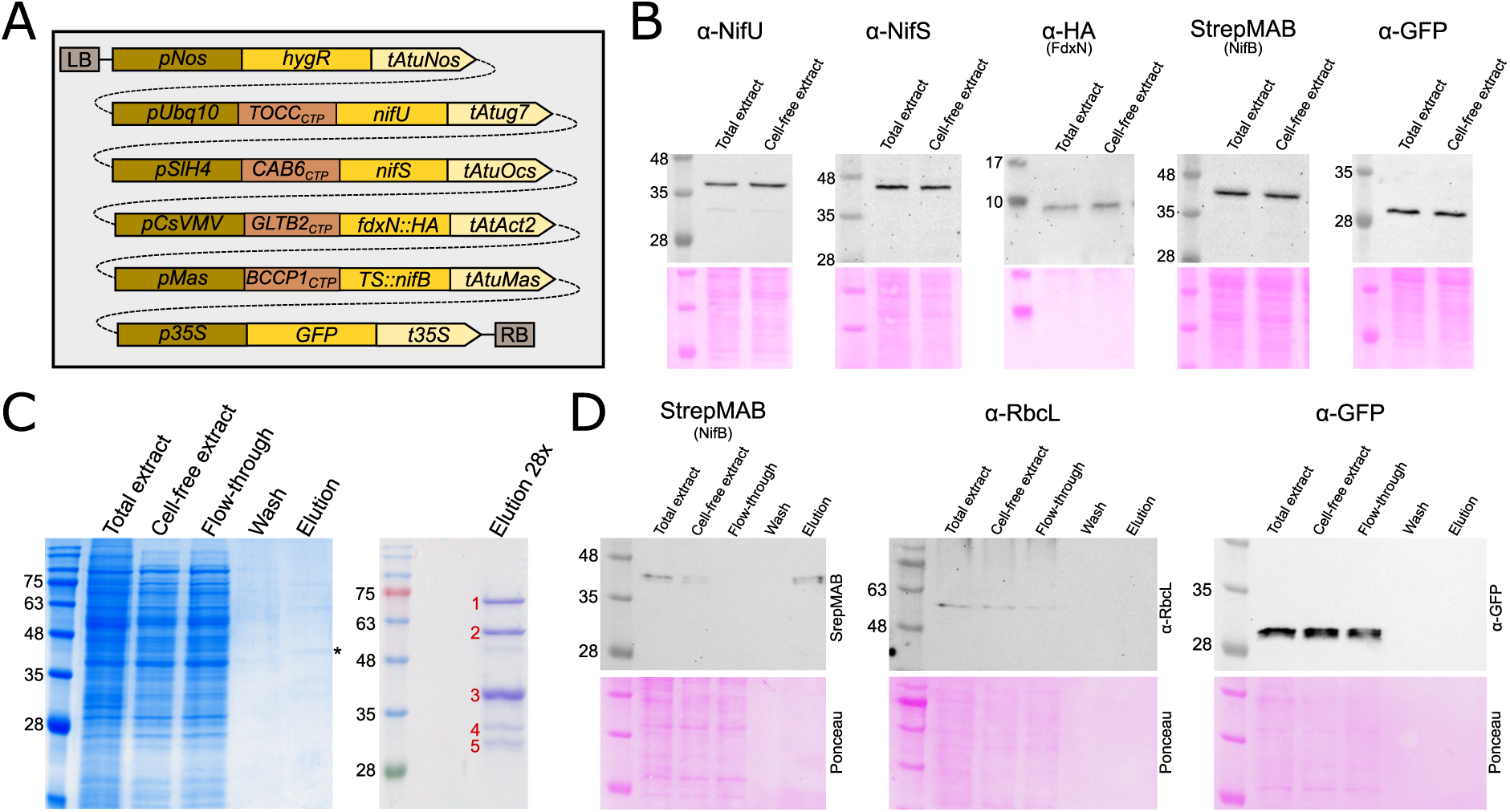
Multi-gene stacking of the NifB module in stably transformed plant cells. (A) Construct design for stable transformation for expression plastid-targeted NifU, NifS, FdxN::HA, Twin-Strep::NifB, and cytosolic GFP. CTP: chloroplast targeting peptide (B) Western blots and ponceau-stained membranes of total and soluble plant extracts from MM1 plant cells transformed with the construct depicted in (A) with *nifB* from *Methanobacterium bryantii*, using α-NifU, α-NifS, α-HA, α-StrepII and α-GFP antibodies. (C) Left panel: Coomassie-stained SDS-PAGE gel loaded with different fractions of the Strep-Tactin affinity chromatographic purification of Col-0 cells transformed with the construct depicted in (A) with *nifB* from *Methanosarcina acetivorans*. Asterisk indicates the expected mass of TS-NifB, 39.5 kDa. Right panel: concentrated elution. Bands labelled with red numbers were analyzed by mass spectrometry. Band 3 was identified as TS::NifB, whereas bands 1, 2, and 4 corresponded to co-purified Arabidopsis proteins methylcrotonoyl-CoA carboxylase subunits alpha (Q42523) and beta (Q9LDD8), and the biotin carboxyl carrier protein of acetyl-CoA carboxylase (F4KE21), respectively. Band 5 was identified as a mix of the Arabidopsis proteins A0A384KVD0 and Q9SSR9 of unknown functions. (D) Western blots of the purification steps shown in (C) with the α-Strep (StrepMAB), α-RbcL, α-GFP antibodies.

## Discussion

In this study, we established and validated a rapid and efficient workflow for transient and stable expression of multigene constructs in *A. thaliana* suspension cultures, with the aim of enabling complex engineering tasks such as the reconstruction of the nitrogen fixation pathway in plant cells. Our approach is based on the plant cell pack infiltration method (Rademacher *et al*., 2019) and adapts it for Arabidopsis, demonstrating robust transformation efficiency, compatibility with several cell lines, and rapid generation of stably transformed cultures. By applying this system to the expression of plastid-targeted Nif proteins, we showcase its utility for synthetic biology applications. A key innovation of this work is the combination of transient and stable transformation workflows into a single streamlined platform. Unlike previous methods, which require more than 40 days to yield 200 ml of transgenic cell cultures (García-León *et al*., 2018) or 29 days to yield 42 ml of culture (Van Leene *et al*., 2011), the workflow presented here can generate 300 ml transgenic culture containing 100 g of cells in less than 2 weeks. This acceleration is probably due to a combination of a higher amount of starting material and a higher transformation efficiency. The workflow relies on basic low-cost equipment found in most biology laboratories. It significantly increases experimental throughput, which is especially important for iterative testing of multi-component systems like nitrogenase biosynthesis. Furthermore, transformed cultures could be propagated without the need for selection, opening the way for simple marker-less transformation applications.

While these features represent major advantages for plant synthetic biology, there are also limitations to consider. The current workflow yields heterogeneous cell populations rather than clonal lines, leading to varying T-DNA integration sites and expression profiles. While this population-based approach suffices for solubility screening and expression optimization, applications requiring precise genotype-phenotype correlations may benefit from purified transgenic lines. However, isolating and propagating individual Arabidopsis suspension cells is technically challenging due to their requirement for high cell densities (Muir *et al*., 1954). This constraint can be addressed through the use “feeder cells” that maintain the growth of isolated transformant cells or through microdroplet-based approaches, in which single cells are cultured in small volumes (Schweiger *et al*., 1987; Spangenberg and Koop, 1992). The ability to isolate purified cell lines will also be critical for forward-genetic screens, which are another possible future application of the presented platform thanks to the high transformation efficiency combined with the possibility to generate stable transformants. For example, bulk-transformations of fully assembled *nif* libraries could be employed to select for nitrogenase activity using fluorescent H_2_ or ethylene biosensors in combination with fluorescence activated cell sorting.

The NifB solubility screen yielded results consistent with previous observations in *Nicotiana benthamiana* leaves, supporting the use of Arabidopsis suspension cultures as a relevant model system for plastid engineering. Specifically, the only three NifB homologs that had been shown to be soluble in *N. benthamiana* leaves when targeted to chloroplasts and/or mitochondria were also included in the library of the present study and all of them were soluble in our Arabidopsis system. Notably, none of the 26 previously tested bacterial NifB homologs were soluble in *N. benthamiana* (Jiang *et al*., 2022). Similarly, in the present study, none of the two tested bacterial NifB homologs were soluble in Arabidopsis cells as opposed to 11 out of 13 tested archaeal homologs that exhibited some level of solubility. This provides further evidence that archaeal Nif variants might generally be superior for engineering BNF in eukaryotes.

This work lays the foundation for implementing multi-component pathways in plastids and for exploring the effects of subcellular localization and nutrient availability on nitrogenase stability and activity. Some Arabidopsis suspension cultures can be readily shifted between heterotrophic and photomixotrophic conditions, enabling controlled studies on the impact of photosynthetically produced oxygen on Nif protein accumulation and function. However, not all cell lines are capable of consistently developing chloroplasts even under light conditions, as shown for Col-0 (Fig. 2C); therefore, the cell line MM1 would be more suitable to address research questions related to chloroplast dynamics. This flexibility, in combination with high transformation speed, makes the system highly suitable for prototyping nitrogenase expression strategies and other complex biosynthetic pathways.

## Abbreviations

2,4-D: 2,4-dichlorophenoxyacetic acid
BNF: biological nitrogen fixation
CTP: chloroplast targeting peptide
MS: Murashige and Skoog
NAA: 1-naphthaleneacetic acid
PCP: plant cell pack
TS: Twin-Strep

## Supplementary data

Figure S1: *nifB* library design and homolog selection criteria.

Figure S2: Assessment of *A. tumefaciens* strain EHA105 and cell line YG1 for PCP infiltration as well as examples of stable transformation of different cell lines.

Figure S3: NifX solubility testing.

Figure S4: Localization controls for confocal microscopy.

Figure S5: Additional replicate of NifB solubility screening.

Figure S6: NifB protein confirmation by mass spectrometry.

Table S1: Primers used in this study.

Table S2: Plasmids used in this study.

Table S3: Synthetic sequences ordered for this study.

Table S4: NifB library and corresponding source organisms.

## Acknowledgements

We thank Álvaro Eseverri for help with MoClo assemblies, David Alabadí for providing the cell line PSB-L, and Laszlo Bako for providing the cell lines Col-0 and MM1.

## Author contributions

L.M., G.A.T., Z.F.E., X.J., and S.B. performed the experimental work, L.M., G.A.T., and L.M.R. designed experiments, analyzed data, and wrote the manuscript.

## Conflict of interest

The authors declare no conflicts of interest.

## Funding

This work was supported, in whole or in part, by the Gates Foundation grants INV-005889 and INV-067006. Under the grant conditions of the Foundation, a Creative Commons Attribution 4.0 License has already been assigned to the Author Accepted Manuscript version that might arise from this submission. L.M. was supported by the Swiss National Science Foundation grant P2EZP3_199998. G.A.T. was supported by Programa Propio UPM de I+D+i 2022.

## Data availability

All data supporting the findings of this study are available within the paper and its Supplementary Information. A preprint version of the manuscript is available at bioRxiv under a CC BY 4.0 license.

## Notes

### Competing Interest Statement

The authors have declared no competing interest.

### Summary of Updates

Supplemental files added. Supplemental Figures S1-S6. Supplemental Tables S1-S4.

