## Supplementary material for "A high-throughput heterologous expression platform for plant synthetic biology based on Arabidopsis suspension cells": Suplemental Figures S1-S6

[2] Present address, Institute of Microbiology, ETH Zurich, 8093 Zurich, Switzerland.

### Supplementary data

Figure S1: *nifB* library design and homolog selection criteria.

Figure S2: Assessment of *A. tumefaciens* strain EHA105 and cell line YG1 for PCP infiltration as well as examples of stable transformation of different cell lines.

Figure S3: NifX solubility testing.

Figure S4: Localization controls for confocal microscopy.

Figure S5: Additional replicate of NifB solubility screening.

Figure S6: NifB protein confirmation by mass spectrometry.

Table S1: Primers used in this study.

Table S2: Plasmids used in this study.

Table S3: Synthetic sequences ordered for this study.

Table S4: NifB library and corresponding source organisms.

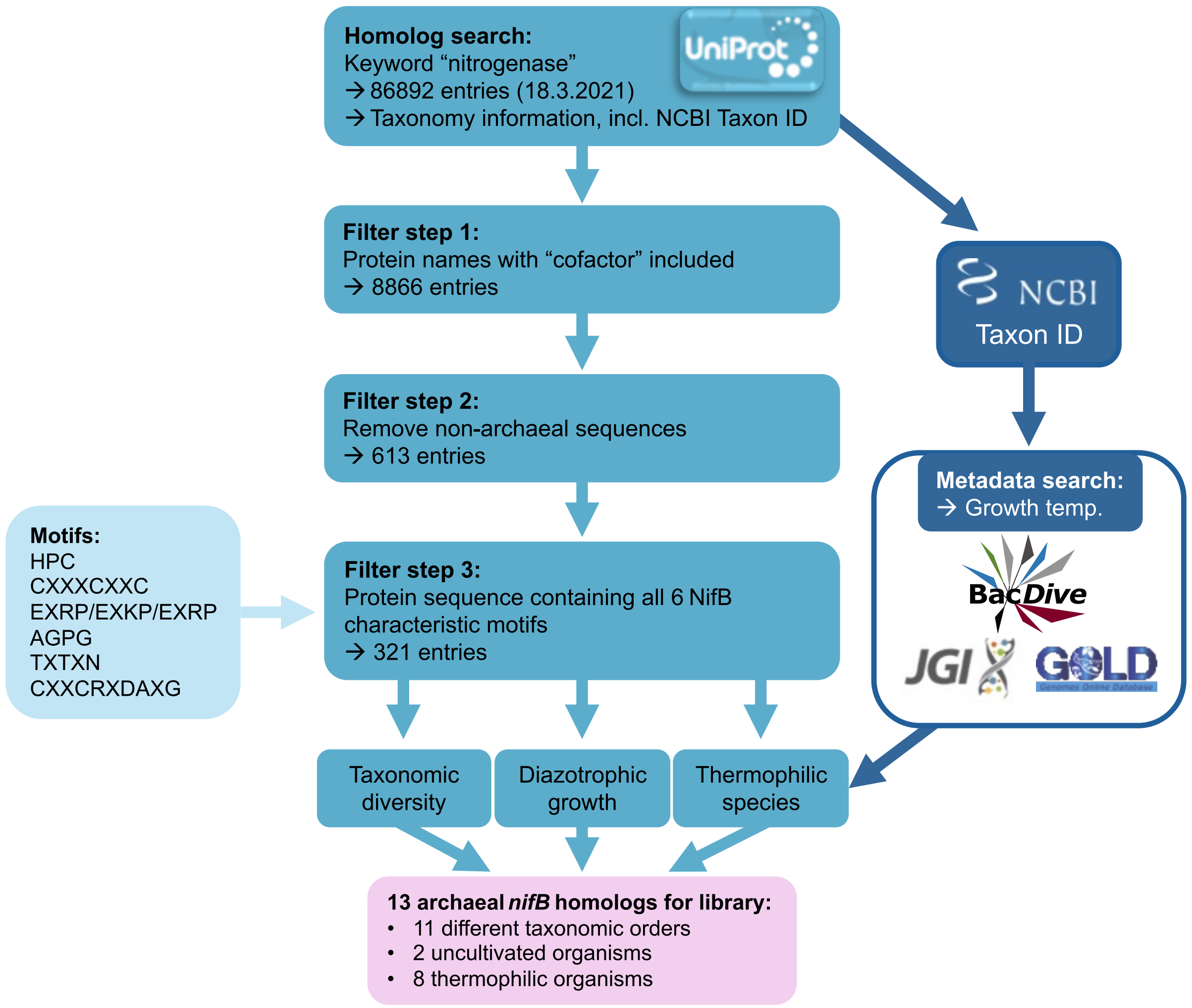


**Fig S1: *nifB* library design.** Archaeal NifB proteins present in the Uniprot database (Bateman *et al.*, 2021) were identified by first obtaining a list of all entries with the keyword “nitrogenase”, then filtering this dataset by presence of the word “cofactor” in the protein name, by archaeal origin, and by the presence of six NifB protein motifs (Arragain *et al.*, 2017). The dataset obtained from Uniprot contained information on the taxonomy of source organisms, which was used to select a diverse selection of NifB sequences, as well as the NCBI Taxon ID, which was used to obtain information on growth temperature, habitats, and metabolism. Main selection criteria were taxonomic diversity of the library, high growth or isolation temperature, extreme habitats, and reported diazotrophy.

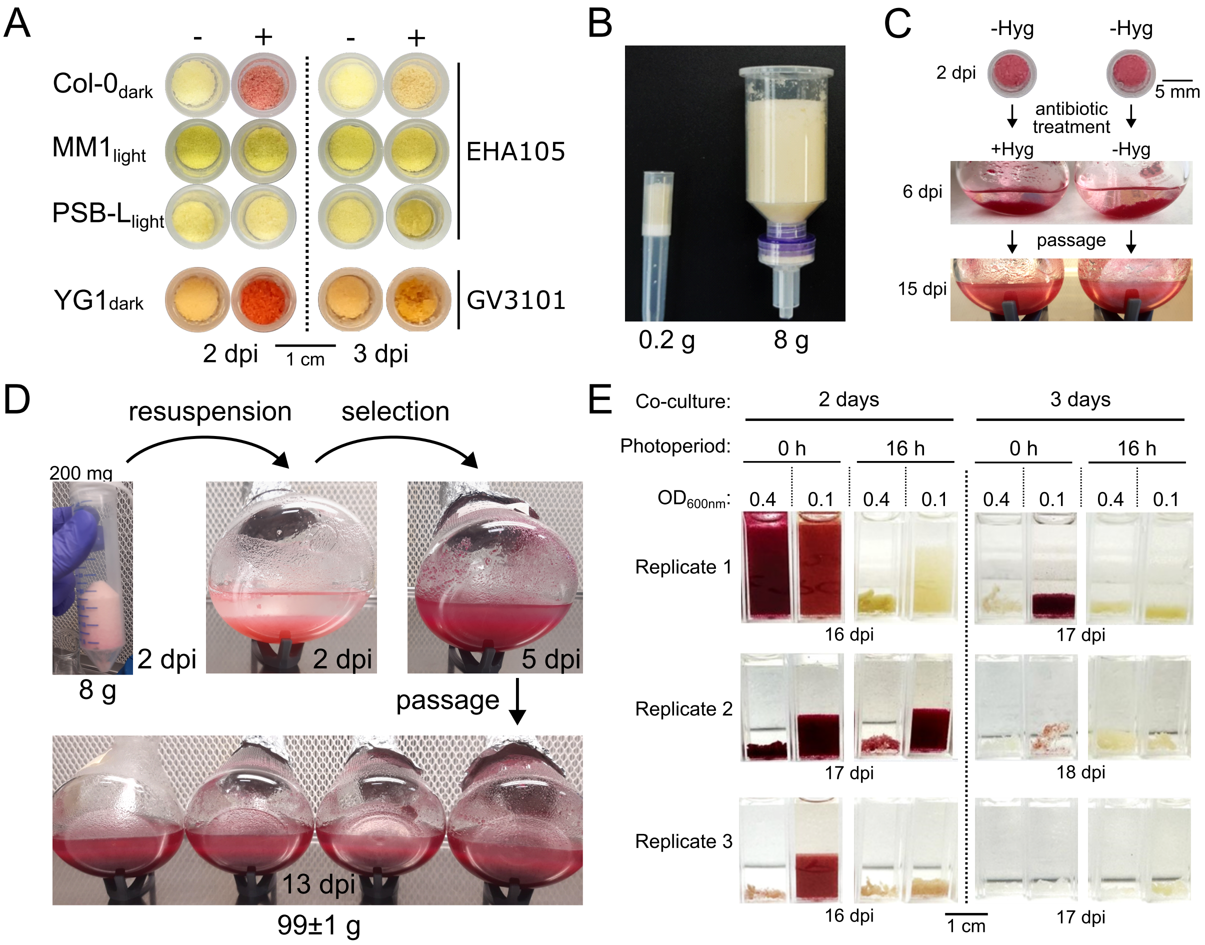


**Fig S2: Plant cell pack (PCP) infiltration is a simple and fast method for transient and stable transformation of Arabidopsis suspension cells**. (A) PCPs cast from 500 µl dense cell suspension and infiltrated with the *A. tumefaciens* strains EHA105 and GV3101::pMP90 harboring a *35S::RUBY* reporter cassette (+) or with infiltration buffer only (-). (B) A 1000-µl filter pipette tip and a midi-prep column with a 15-ml reservoir loaded with PCPs before infiltration. (C) 500-µl PCPs of cell line Col-0 infiltrated with the *35S:RUBY* reporter cassette 2 days after infiltration and after resuspension in growth medium and a culture passage with and without hygromycin selection. (D) Different stages of biomass production of stably transformed suspension cells derived from an 8-g PCP. Shown are a PCP in a 50-ml centrifugation tube, cells after resuspension, after a 3-day selection with hygromycin, and after a passage from one to four flasks. (E) Recovery of stable transformants of MM1 cells after treatment of 2 or 3 days with an *A. tumefaciens* strain carrying *35S:RUBY* at different optical densities and selected with hygromycin for 1 week. Pictures show cultures after transferring to a spectrophotometry cuvette and sedimentation. Before PCP infiltration, suspension cultures were grown either under a 16-h photoperiod or without light. Shown are 3 biological replicates. OD values refer to the *A. tumefaciens* suspensions used for PCP infiltration. dpi: days post infiltration.

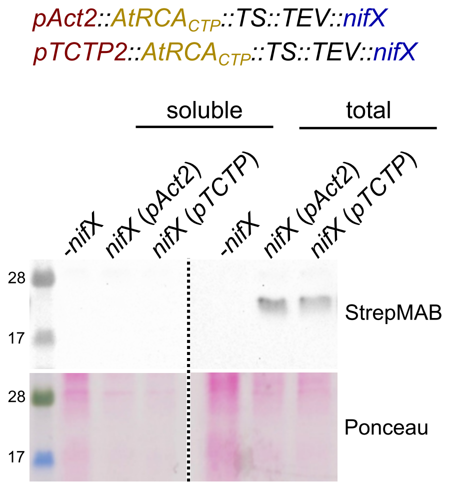


**Fig S3: NifX is expressed but insoluble in plant cell packs.** Western blots and ponceau-stained membranes of total and soluble protein extracts of plant cell packs transformed with *nifX* driven by the Arabidopsis *act2* or *TCTP* promoters. The protein was fused to the minimal version of the Arabidopsis RCA chloroplast targeting peptide (Eseverri *et al.*, 2020a), to the TS-tag for detection, and to the TEV protease cleavage site.

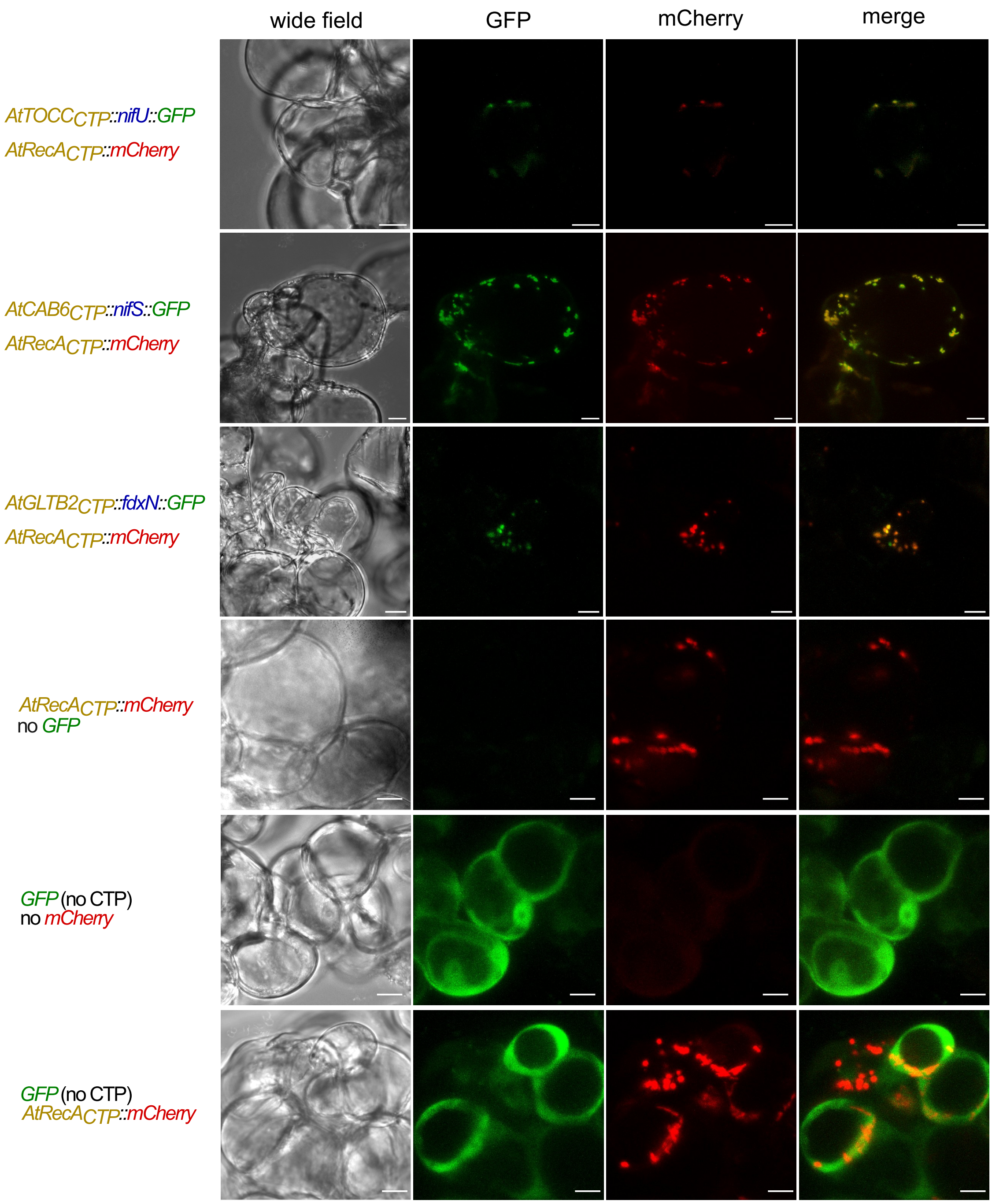


**Fig S4: Subcellular localization of plastid-targeted Nif proteins.** Detection of GFP signal in Arabidopsis cells transformed with *nifU*, *nifS* and *fdxN* fused to *gfp* and the TOCC, CAB6 and GLTB2 CTPs, respectively, by confocal laser scanning microscopy (rows 1-3). Arabidopsis suspension cells expressing localization control constructs are also shown. Row 4: mCherry fused to the Arabidopsis RecA chloroplast targeting peptide. Row 5: GFP lacking a chloroplast targeting peptide, showing diffuse cytosolic localization. Row 6: Co-expression of both controls from a single plasmid. For each construct, representative wide-field images and individual GFP and mCherry channels, as well as merged fluorescence channels are shown. Scale bars: 10 µm.

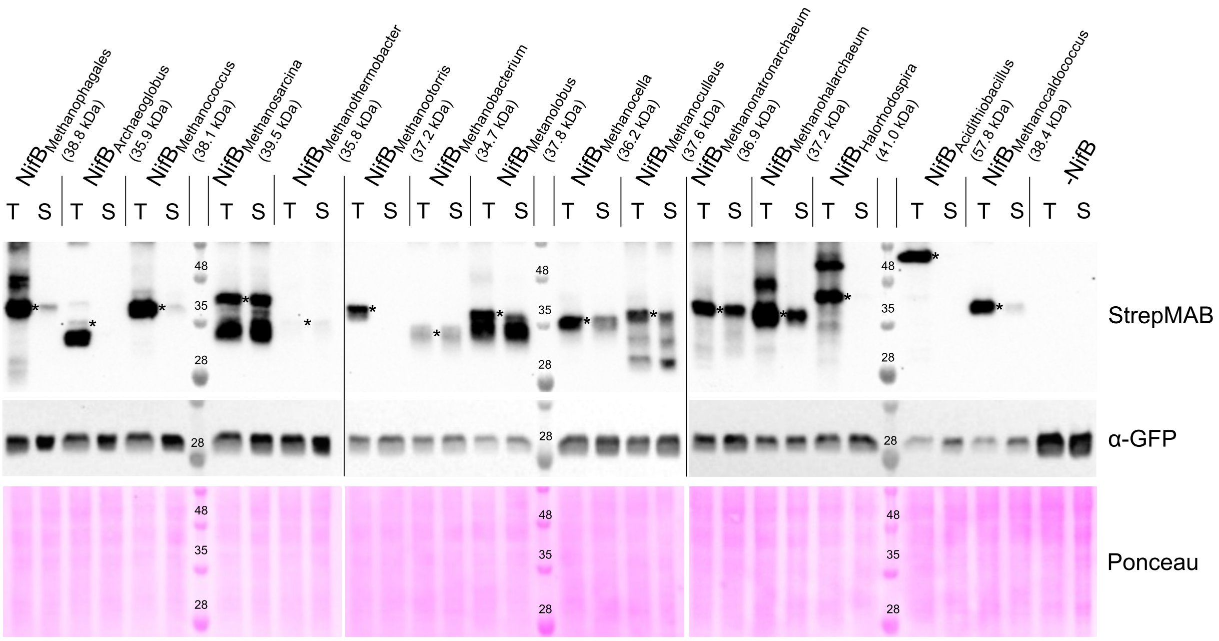


**Fig S5: Solubility screening identified new archaeal NifB protein variants suitable for plant expression.** Replicate of experiment shown in Fig. 5. Western blots and ponceau-stained membranes of total (T) and soluble (S) plant cell extracts. NifB variants were fused to the Twin-Strep tag for detection with a Strep monoclonal antibody (StrepMAB). A control with the same expression vector but without *nifB* was included (-NifB). Asterisks mark the positions of full-length NifB proteins.

1 MASSSFSVTS PAAAASVYAV TQTSSHFPIQ NRSRRVSFRL SAKPKLRFLS

51 KPSRSSYPVV KAASAWSHPQ FEK**GGGSGGG** **SGGSAWSHPQ** **FEKSSMPEEN**

101 **QPIKEKNNGP** **ILGEELLR**KI SEHPCYDKNA QHKYGR**IHLA** **VAPACNIQCN**

151 **FCVREFDCVN** **ESRPGVTSKV** **LTPEEALEK**T K**QILAEYPFI** **KVVAIAGPGD**

201 **PLANDETFET** **FELIRNEFPE** **ITLCMSTNGL** **MLPEKLPEIL** **RTGVSTLTVT**

251 **VNAIDPEIQA** **KIVDHIFYHG** **K**VYKGVEAAK IQIKNQLDGI KAAIDAGIVV

301 K**VNTVLIPGI** **NDKHIIEIAK** K**LNELGVYIM** **NVMPLINQGA** **FADLEPPTPE**

351 **ERKAVQEACE** **PYVMQMR**HCR QCR**ADAYGLL** **AQDMSQMSEE** **RR**KVIKIQTK

401 EDMEKARAVL EKNGKKEA

**Fig S6: Peptide mass fingerprinting of purified TS::NifB protein.** Sequence of $BCCP1_{CTP}$::TS::NifB of *Methanosarcina acetivorans*. Blue residues show the part of the BCCP1 chloroplast targeting peptide predicted to be cleaved during transfer to the plastid (Eseverri *et al.*, 2020b). Red bold residues show the peptides identified by mass spectrometry.
